# Revealing Atomic-scale Molecular Diffusion of a Plant Transcription Factor WRKY domain protein along DNA

**DOI:** 10.1101/2020.02.14.950295

**Authors:** Liqiang Dai, Yongping Xu, Zhenwei Du, Xiao-dong Su, Jin Yu

## Abstract

Transcription factor (TF) target search on genome is highly essential for gene expression and regulation. High-resolution determination of TF diffusion along DNA remains technically challenging. Here we constructed a TF model system of the plant WRKY domain protein in complex with DNA from crystallography and demonstrated microsecond diffusion dynamics of WRKY on the DNA employing all-atom molecular dynamics (MD) simulations. Notably, we found that WRKY preferentially binds to the Crick strand of DNA with significantly stronger energetic association than to the Watson strand. The preferential binding becomes highly prominent from non-specific to specific DNA binding, but less distinct from static binding to diffusive movements of WRKY on the DNA. Remarkably, without employing acceleration forces or bias, we captured a complete one-base pair (bp) stepping cycle of WRKY tracking along major groove of DNA with homogenous (AT)_n_ sequence, as individual protein-DNA contacts break and reform at the binding interface. Continuous tracking of WRKY forward or backward, with occasional sliding as well as strand crossing to the minor groove of DNA, have also been captured in the simulation. The processive diffusion of WRKY had been confirmed by accompanied single-molecule fluorescence assays and coarse-grained (CG) structural simulations. The study thus provides unprecedented structural dynamics details on the TF diffusion, suggests how TF possibly approaches to gene target, and supports further high-precision experimental follow-up. The stochastic movements revealed in the TF diffusion also provide general clues on how other nucleic acid walkers step and slide along DNA.

**Significance Statement:** How transcription factors search for target genes impact on how quickly and accurately the genes are transcribed and expressed. To locate target sufficiently fast, 1D diffusion of the protein along DNA appears essential. Experimentally, it remains challenging to determine diffusional steps of protein on DNA. Here, we report all-atom equilibrium simulations of a WRKY protein binding and diffusing on DNA, revealing structural dynamics details which have not been identified previously. We unprecedently demonstrate a complete stepping cycle of the protein for one base pair on DNA within microseconds, along with stochastic stepping or sliding, directional switching, and strand crossing. Additionally, we have found preferential DNA strand association of WRKY. These suggest how protein factors approach toward target DNA sequences.

## Introduction

The search and recognition processes of Transcription factors (TFs) on DNA are of fundamental importance in gene expression and regulation. To locate sufficiently fast a target site on genome that is wrapped within three dimensional cellular space, the TFs may proceed with a facilitated diffusion process, alternating between one dimensional (1-D) movements along DNA and three dimensional (3-D) intra-cellular diffusion (1-6). Experimental detection on protein searching motions or 1-D diffusion along DNA have provided evidence on the facilitated diffusion (7-12). Nevertheless, as protein movements for base pair (bp) distances on DNA can take place as fast as microseconds, tracking the 1-D protein diffusion at such a high temporal and spatial resolution remains technically challenging (13-16).

On the other hand, high resolution determinations of protein-DNA complex structures (17) allow one to investigate corresponding conformational dynamics by employing all-atom molecular dynamics (MD) simulations, via high-performance computing (18-20). The protein recognition on specific DNA has been actively examined in recent years using the MD technologies (21-25). In comparison, the protein with non-specific DNA has been less examined. It is commonly expected that nonspecific association and movements of protein on the DNA happen quite slowly and cannot be well sampled via the atomistic MD. Indeed, either comparatively short MD simulations (nano or sub-microseconds) were conducted (21), or external forces were added to accelerate the protein movements or enhance samplings, such as by employing targeted MD or umbrella sampling simulations (23, 26-28). In case that comparatively long or extensive MD simulations have been conducted, one recent study concentrates on association processes of a chromatin protein with DNA(29), but not yet the protein movements. For exemplary all-atom simulation studies on the protein movements along DNA, however, the proteins of concerns are motor proteins such as RNA polymerases(30, 31), or the single-stranded DNA-binding protein(32). In this work, we focus on a model TF and present all-atom microseconds equilibrium simulations of the diffusion dynamics of the TF protein along the double stranded (ds) DNA. The protein factor under our current investigation is a WRKY domain protein from *Arabidopsis thaliana* WRKY1.

WRKY proteins are a large family of transcription factors (TFs) in plants playing important functions for a broad range of signal response, stress control, and disease resistance (33, 34). The number of WRKY family members in *Arabidopsis* reaches over 70, and all of them include a DNA binding domain about 60 amino acids that is called the WRKY domain. The WRKY domain proteins are featured by a highly conserved ‘WRKYGQK’ sequence and a zinc finger motif, both of which turn out to be indispensable for maintaining the DNA binding function. Previously, an *apo* C-terminal domain structure of *Arabidopsis* WRKY1 had been made available (35). Recently, a high-resolution crystal structure of the N-terminal WRKY domain protein in complex with a specific DNA binding sequence is obtained (36). Based on this structure, we performed atomistic MD simulations on the protein-DNA complexes (with a 34-bp dsDNA) in explicit solvent conditions, constructed both for the specific and non-specific DNA binding systems. We identified comparatively strong association of WRKY with the Crick (sense) strand, and weak association with the other (Watson or anti-sense) strand, which demonstrates most prominently in the specific DNA binding case. Notably, our simulations further captured 1-bp cyclic stepping motions for a full set of protein residues forming contacts on the DNA, as the protein track along the DNA groove forward or backward, spontaneously. Moreover, the simulations have also revealed protein sliding at a larger stepping size (> 1 bp) or more stochastically crossing the DNA strand. The processive diffusion of the WRKY domain protein along DNA have been confirmed by accompanied single-molecule fluorescence assays and coarse-grained (CG) simulations in this work.

## Results

### Specific to non-specific DNA association of WRKY and an onset of diffusion

We conducted microseconds equilibrium MD simulations on the WRKY-DNA complexes, with a specific binding motif and a non-specific DNA sequence, respectively. The specific complex had been constructed directly from the crystal structure obtained (36) (see **SI Methods**), while the non-specific complex structure was modeled by converting the specific core sequence of DNA (CT<u>GGTC</u>AAAG) in the crystal structure to a nonspecific one (CT<u>GATA</u>AAAG) (see **SI Methods**). Using the isothermal titration calorimetry (ITC), we had determined the WRKY dissociation constants with DNA on the above specific and non-specific sequences as K_D_=0.1 μM and 8 μM, respectively (see supplementary **Fig S1**).

By conducting and comparing two 10-μs MD simulations of WRKY on the specific and non-specific sequences (see **Fig 1**), one notices a well localization of WRKY on DNA around the specific motif, with a restricted amount of longitudinal (ΔX ∼ 0.2± 0.8Å) and rotational movements (ΔΘ∼ −0.9±5.7°) of the protein center of mass (COM) after a 2-μs pre-equilibration period. In comparison, WRKY modeled on the non-specific DNA sequence demonstrated a positional relaxation or an onset of diffusion along DNA by shifting soon both longitudinally and rotationally (ΔX ∼2.8±1.4 Å and ΔΘ ∼ 28.1±6.1°). Indeed, the WRKY domain protein tracks slightly along the major groove in the simulation (with an ionic concentration set at 150 mM). Structural alignments according to the associated DNA segment also suggest that conformation changes of the non-specific complex (ΔRMSD ∼ 8 Å) are substantially larger than that of the specific complex (ΔRMSD ∼ 3 Å). Two movies are provided for viewing the specific and non-specific DNA binding complexes of WRKY in the simulation (see **SI Movie S1** and **S2**), respectively.

**Fig 1.**
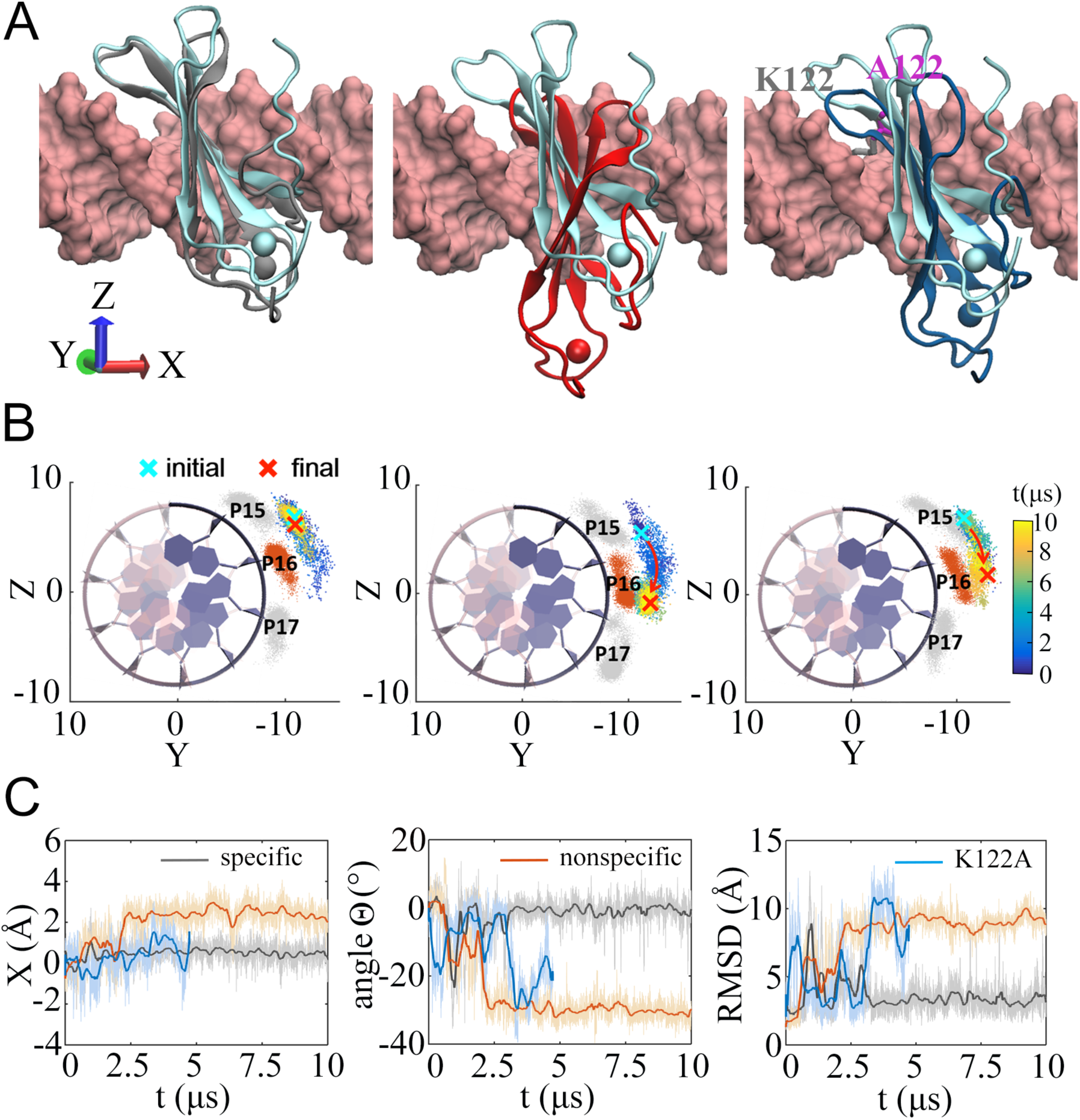
Specific and non-specific DNA association of WRKY. (A) Comparisons of the initial (cyan) and final (gray, red, blue) structures of the simulation of the wild-type (wt) protein binding on the specific DNA *(left)*, the non-specific DNA *(middle)*, and the mutant (mt) protein (K122A) protein binding on the original specific DNA *(right)*. The XYZ-axis is denoted and the protein moves along DNA longitudinally following the X direction. (B) The rotation of the center of mass (COM) of the protein along DNA projected onto the Y-Z plane. The initial and final positioning of the protein COM are labeled. The time evolution is represented by changing colors (from blue to yellow). The COMs of the phosphates of the protein associated nucleotides are also shown in gray (P15 and 17) or orange (P16) clouds. (C) The longitudinal movements X, rotation angles Θ, and the RMSDs of the simulated protein-DNA complexes, for respective simulation systems (wt specific for 10-μs, dark; non-specific for 10-μs, orange; and K122A mt complexes for 5-μs, blue).

Meanwhile, we also constructed a mutant (mt) WRKY K122A with a lowered DNA affinity: K_D_ ∼ 1 μM (see **SI Methods** and **SI Fig S1**). Correspondingly, we performed MD simulation for this mutant complex, modeled on the original ‘specific’ DNA sequence. The results show that the mt-WRKY started shifting along DNA similarly as the non-specific wild-type (wt) complex (also see **Fig 1**). The measurements on the positional and structural deviations of the K122A mutant also show intermediate behaviors in between the specific and the non-specific ones.

### WRKY association with DNA is strongly biased on one strand and the bias is most stably maintained in in the specific binding case

By close examinations, we identified detailed interactions at the protein-DNA interface, for both the specific and the non-specific binding systems (see **Fig 2**). In particular, we found substantial hydrogen bonding (HB) interactions between the WRKY domain protein and the Crick strand DNA (e.g. 7-10 HBs), in both specific and non-specific cases. In the specific binding (**Fig 2A** *bottom left*): Y119(O)-C18(N4), Y119(OH)-T17(O2P), K122(NZ)-G16(O6), K125(N)-G15(O2P), R131(NH1)-G16(O2P), Y133(OH)-G16(O2P), R135(NH1/NH2)-C18(O2P), K144(NZ)-T17(O1P), and Q146(NE2)-G16(O1P), while Q146(NE2) also has a water mediated HB with T17(O2P). Among them, some charged residues such as arginine or lysine (R131, R135, and K144) also form electrostatic or salt-bridge interactions with the negatively charged phosphate groups on the DNA (see **SI Fig S2A**); some residues are polar (Y119, Y133, and Q146) and form HBs with the DNA backbone; Y119 additionally forms a HB with the C18 base, while K122 also forms a HB with the G16 base. Hence, the very specific recognition to the core sequence seems to be achieved mainly by K122 and Y119. In contrast, there are much fewer interactions formed between WRKY and the other DNA strand, the Watson strand (e.g. ∼ 2 HBs, plus 3 water-mediated HBs addressed later): It mainly involves HB or salt-bridge interaction from the charged R117 and K118 with the DNA backbone. The schematics summarizing the HBs and salt-bridges between the WRKY domain protein and specific DNA strands are found in **SI Fig S2A**.

**Fig 2.**
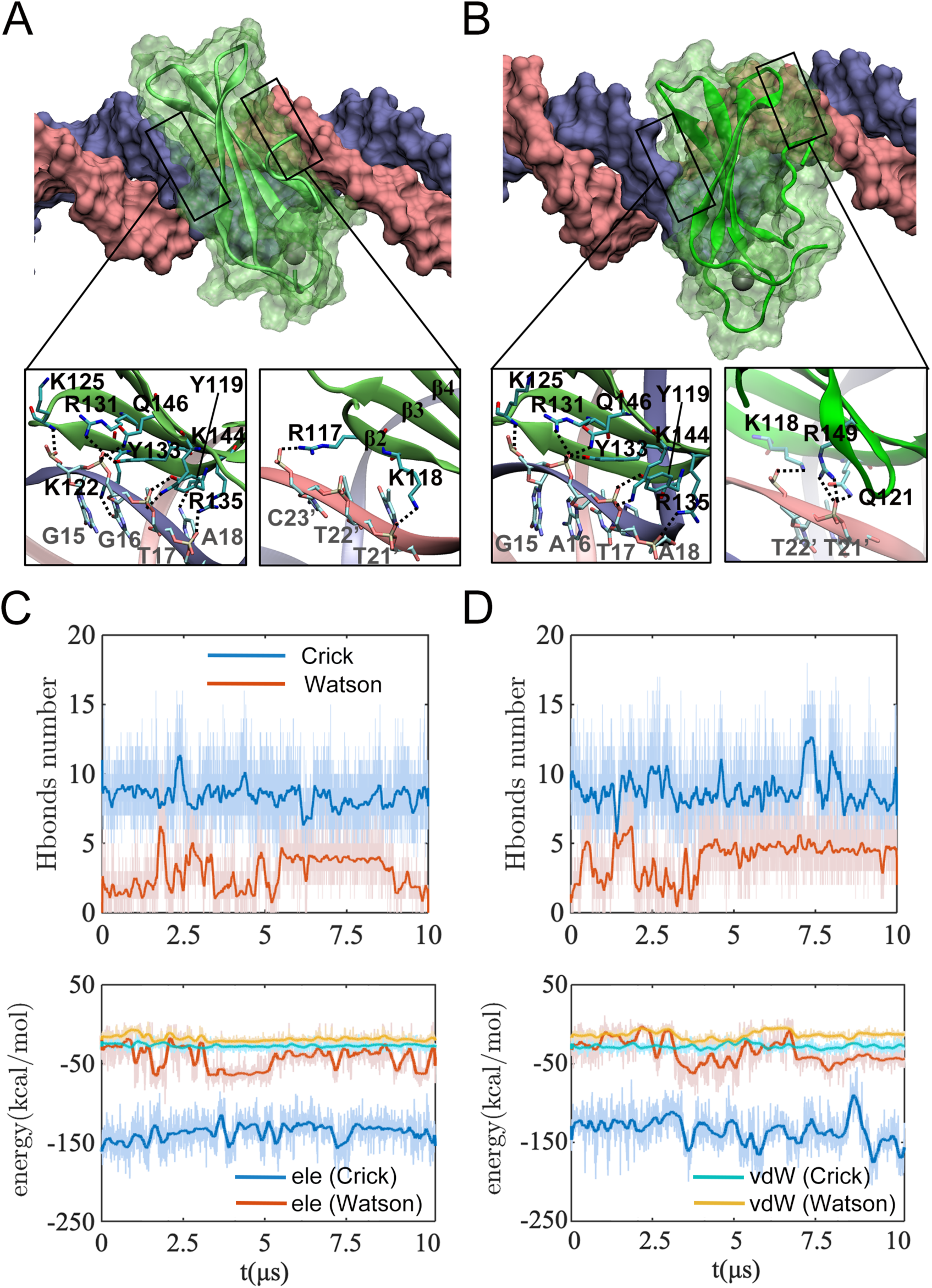
The association between the WRKY domain protein and respective strands of DNA. (A) WRKY on the specific DNA (core sequence: GGTC) and (B) the non-specific DNA (core sequence replaced by: GATA). The structures of the respective protein-DNA complexes are shown toward the end of the 10-μs simulations (*top*). The protein is colored in green, and the DNA strands are shown in blue (the Crick strand) and pink (the Watson strand). The hydrogen bonding (HB) associations at the protein-DNA interface are shown in molecular details (*bottom*). The HB interaction is defined by a cut-off distance of 3.5 Å between the donor and acceptor atoms and an associated donor-hydrogen-acceptor angle of 140° (HBs formed >5% the simulation time are shown). The numbers of HBs counted dynamically during simulation are also shown (C) and (D), for the specific and non-specific binding cases, respectively; the electrostatic (ele) and van der Waals (vdW) interactions at the binding interface between the protein and the Crick (blue, cyan)/Watson (red, orange) strand are calculated (C): ele (−137±17 /-43±19 kcal/mol) and vdW (−27±4 / −18±5 kcal/mol); (D) ele (−135±22 / −35±19 kcal/mol) and vdW (−28±5 / −13±6 kcal/mol).

In comparison, one can see that the HB association of WRKY to the non-specific DNA include (**Fig 2B** *bottom*): Y119(OH)-T17(O2P), K125(N)-G15(O2P), R131(NH1)-A16(O2P), Y133(OH)-A16(O2P), R135(NH1/NH2)-A18(O2P), K144(NZ)-T17(O1P), and Q146(NE2)-A16(O1P) as to the Crick strand; Q22(NE2)-T21’(O2P), R50(NH1)-T22’(O2P), G54(N)-T22’(O2P), and Q55(NE2)-C23’(O2P) as to the Watson strand. In comparison with the specific DNA case, the Crick strand HB association now demonstrates slightly larger fluctuations. The highly ‘specific’ K122 association with the core sequence base (on the Crick strand) becomes missing. The WRKY association with the other strand (the Watson strand) appears to strengthen by an increasing number of HBs from the specific to the non-specific case (see **Fig 2C** and **D** *top*), while energetically it does strengthen slightly as well (after ∼ 4 μs). Indeed, the variation of the WRKY association with the Watson strand is due to shifting of the interface between WRKY and DNA, i.e., from the beta strand 2 in the specific binding case to the beta 4&5 strands in the non-specific case (**Fig 2A** and **B** *bottom*). The interaction schematics between WRKY and the non-specific DNA strands can also be found in **SI Fig S2B**.

Accordingly, we calculated the electrostatic (ele) and van der Waals (vdW) interaction energies at the protein-DNA interface, for respective DNA strands and the protein (exclude the flexible terminal region, see **SI Methods**), in both the specific and non-specific binding cases. For both cases (specific & non-specific), the interactions between the Crick strand and WRKY turn out to be similarly strong (see **Fig 2 C** and **D** *bottom*): ele (−137±17 and −135±22 kcal/mol) and vdW (−27±5 and −28±5 kcal/mol), with slightly higher fluctuations for the ele component in the non-specific binding. In comparison, the WRKY interactions with the Watson strand are significantly weaker than with the Crick strand. The Watson strand interaction in the specific binding (ele: −43±19 and vdW −18±5 kcal/mol) still appear stronger than that in the non-specific binding (ele: −35±19 kcal/mol and vdW −13±6 kcal/mol). Indeed, in the specific binding to the Watson strand, besides the two HBs and salt-bridge interactions from R117 and K118, there are three water mediated HBs linking W116, Q121 and Y134 to the Watson strand (W116(O)-water-C22’(O1P), Q121(NE2)-water-T20’(O1P), Y134(OH)-water-T21’(O2P); see **SI Fig S3A**). In contrast, WRKY binds non-specifically to the Watson strand via a loop region in between beta strand 4 and 5, different from that in the specific binding via strand 2, which may account for the instabilities.

In addition, we measured disparities and correlations between the WRKY protein interactions with the Crick strand and that with the Watson strand during simulations, by calculating the t-values and Pearson correlation coefficients between the time series of respective interaction strengths (see **SI Methods**). For the electrostatic association, the calculations show slightly higher t-values and lowers correlation strengths in the WRKY binding with the specific DNA (t-value 182 and correlation −0.07) than with the non-specific DNA (174 and 0.12; see **SI Table S1**), indicating a slightly stronger strand association bias and less correlation between the two-strand associations in the specific DNA binding case.

The hydrophobic interactions have also been monitored in the simulation. The involved hydrophobic residues (see **SI Methods**) with the Crick stand also appear more than that with the Watson strand (see **SI Fig S3B** and **C**), in both the specific and no-specific cases. In addition, we counted numbers of water molecules around the protein (within 5 Å) or at the protein-DNA interface (within 5 Å): Similar amount (slightly more) waters surrounding WRKY in the specific (313 ±6) and non-specific case (308±12), yet a bit less amount of waters at the protein-DNA interface can be identified in the specific binding (∼ 41± 6) than in the non-specific case (∼ 47 ± 8) (see **SI Fig S3D**).

For the mutant(mt) WRKY K122A in complex with the specific DNA, the association with the Crick strand becomes slightly destabilized and more volatile (ele −131±25 and vdW −32±8 kcal/mol) than the specific binding case, likely due to loss of interaction from K122. The mt-WRKY association with the Watson strand (ele −40±26 and vdW −18±7 kcal/mol) is still similar on average to that in the specific binding, while the fluctuations in the electrostatic association enhances significantly. One could indeed see that the binding interface switches close to the loop region in between beta strand 4 and 5, e.g., involving R149 on beta strand 5 as in the above non-specific binding case (see **SI Fig S4**). The ITC measurements of a reduced K_D_ (1μM) comparing to the specific binding case (0.1μM) consistently suggest a lowered DNA affinity of the mt-WRKY. The t-value and Pearson correlation coefficient measured during the simulation between the protein electrostatic associations with the two strands are 126 and −0.12, indicating a lowered disparity or an increased similarity between the two DNA strand associations, comparing with the specific and non-specific binding cases (see **SI Table S1**).

### Atomistic simulation of WRKY diffusion along homogenous DNA sequence (AT)_n_

In the above 10 μs simulation of wt-WRKY dynamics on the non-specific DNA, we have not yet detected diffusion of the protein along DNA: Though overall the protein COM demonstrates longitudinal and rotational motions as for an onset of diffusion, almost all the protein-DNA contacts maintain throughout the simulation (see **Fig 1C** and **SI Fig S2B**). In order to reveal essential diffusional motions of the protein, i.e., the displacements of all the protein-DNA contacts along DNA, we modeled WRKY in complex with homogenous DNA sequences (AT)_n_ (n=30), anticipating that the individual contacts on the DNA would have similar strength so that to allow fast or nearly synchronized residue movements at the protein-DNA interface. We accordingly captured one complete cycle of the diffusion of WRKY, i.e., for 1-bp distance on the DNA, via our atomistic simulation (see **Fig 3)**.

**Fig 3.**
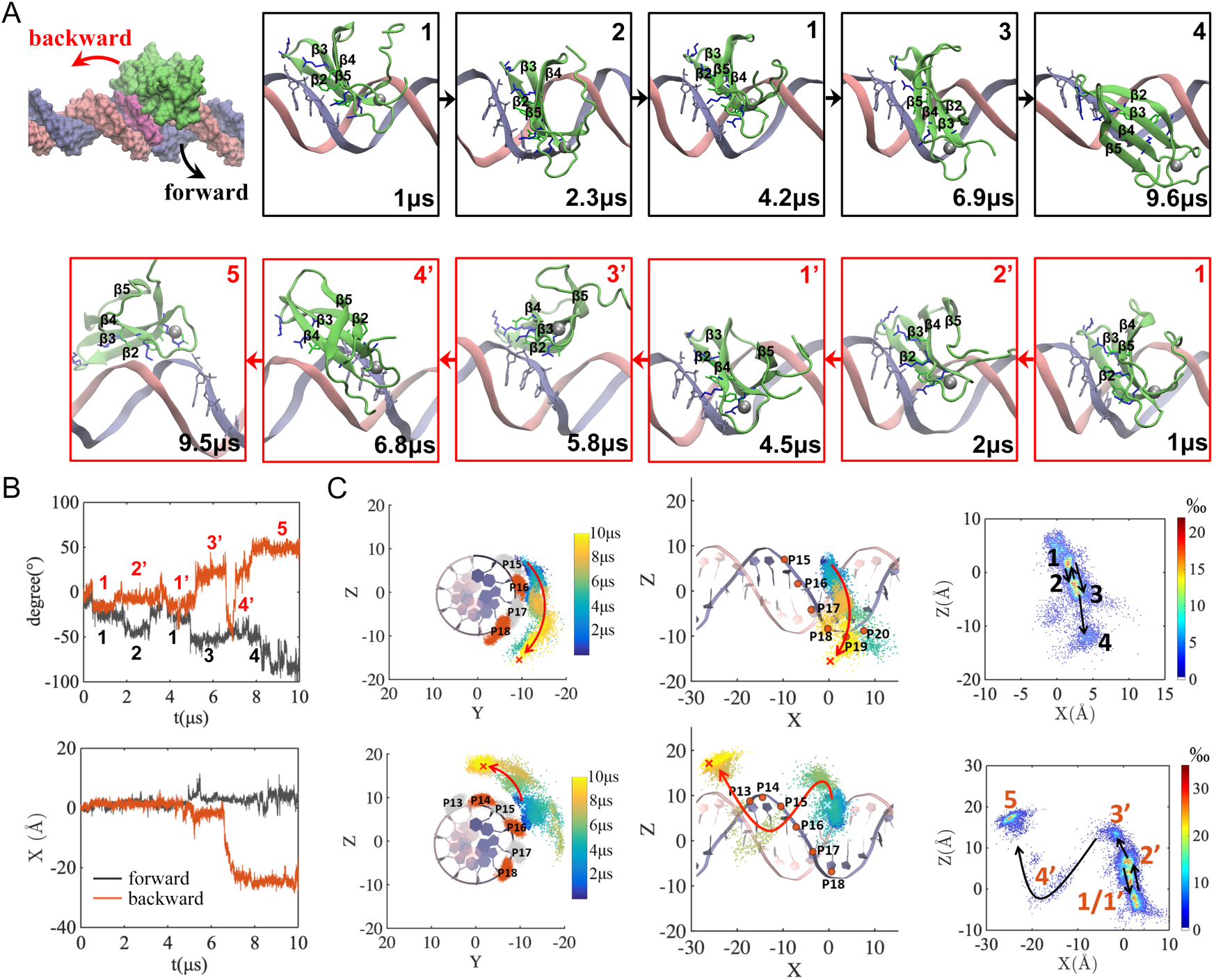
The diffusion of WRKY along DNA in the forward and backward direction revealed from two 10-μs atomistic MD simulations. (A) The representative snapshots taken from the MD trajectories forward (upper, from the left to the right, via state 1 → 2 → 1→3 →4 according to the protein COM movements) and backward (lower, from the right to the left, via state 1 → 2’ → 1’→ 3’ →4’→5, with primed labels to differentiate from the forward states), with the WRKY domain protein shown in green and two DNA strands in blue (the Crick strand) and pink (the Watson strand). (B) The helical trajectories of the protein COM along the DNA, shown for the rotation Θ(t) (upper) and the longitudinal movement X(t) (lower) from the simulation. The coordinate system is defined the same as in Fig 1. Four (forward, dark line) or five (backward, orange line) states are identified along the helical trajectories. (C) The protein COM helical motions along DNA are mapped on the Y-Z plane (left) and then along the Z-X side (middle), colored with the simulation time (blue to yellow). The dsDNA schematics is also shown for an easy view (left: fluctuations of the phosphates are shown in orange and gray as in Fig 1). On the Z-X plane, clustering of the protein COMs into different spatial states are also shown (colored by densities of populations).

In **Fig 3A**, representative snapshots from two MD trajectories are shown, demonstrating WRKY moving forward (along +X direction, or toward right in the figure) and backward (toward left) along DNA, respectively. The longitudinal (along the X axis) and helical circular motions (mapped on the Y-Z plane) of the protein COM along DNA are demonstrated in **Fig 3B** and **C**. In the forward direction, approximately four spatial states reveal, according to relative motions of the protein COM on the DNA: In the first 2 μs, WRKY tracks slightly forward along the major groove of the DNA, closely grabbing on and following the Crick strand (blue), moving from state 1 to 2 (Δx∼1.3 Å and ΔΘ ∼ 18.0°); during 2-5 μs, however, the protein slightly retracks back to state 1; at ∼ 5 μs, the protein quickly steps forward, advancing about 1-bp distance within 0.1-0.2 μs (1→3 transition Δx∼2.5 Å and ΔΘ ∼ 26.2°); after that (> 7.8 μs), WRKY slides forward (3→4 transition Δx∼0.4 Å and ΔΘ ∼15.3°) to adjust its spatial orientation to better align with the major groove at the new site (see **SI movie S3** for viewing the full MD trajectory).

The diffusion captured in the backward direction also starts with some slight forward motions (1→2’) within the first 2 μs, which is then followed by retracking (back to 1), similar to that in the forward trajectory. However, after that, at ∼5 μs, there is a sudden jump of the protein backward (Δx∼2.7Å and ΔΘ ∼ 29.3° for 1→3’). Interestingly, at ∼ 7 μs, we observe a strand-crossing event of WRKY on the DNA in the backward direction. Right after that, WRKY binds onto the minor rather than the major groove of DNA (see **SI Movie S4** for viewing the full MD trajectory).

Since one could not sample processive diffusion of the protein on DNA using the all-atom MD simulations, we conducted coarse-grained (CG) structural dynamics simulations to the WRKY-DNA complex using CafeMol (37). In the CG presentation, each amino acid is represented by a sphere, while each nucleotide is represented by three spheres(38). Correspondingly, there is no specific or the HB type of interactions between protein and DNA. Nevertheless, the electrostatic association between the charged amino acids and phosphate groups on DNA have been taken into account in the CG modeling. In **SI Fig S5**, we show that in the CG representation, WRKY conducts processive diffusion along DNA at variable ionic conditions. In particular, at an ionic strength of 150 mM comparable to a physiological condition (as set in the all-atom MD simulation), WRKY demonstrates mainly the groove tracking behaviors, with occasional strand crossing motions (see **SI Movie S5**), consistent with our observations in the all-atom MD. In comparison, at lower and higher ionic conditions (50 mM and 200 mM), regular groove-tracking motions and frequent ‘micro-dissociation’ events show, respectively (see **SI Movie S6** and **S7**), which correspond well to weak and strong charge screening situations.

In addition, we verified processive diffusion of WRKY on DNA experimentally, using single molecule fluorescence methods (see **SI Methods** and **SI Fig S6**), albeit the resolution is not high enough to discern single bp movements of the protein. The diffusion coefficient has been measured at an order of magnitude of 0.04 bp^2^/μs (see **SI Fig S6D**), which is compatible with our observation of the diffusive behaviors in the microseconds (all-atom) to tens of microseconds (CG) simulations.

### The major groove tracking and residues stepping for one bp cycle of WRKY on the DNA

By close inspections on how individual amino acid residues of WRKY break and reform contacts with DNA during the diffusion, we show detailed stepping dynamics in the forward direction as WRKY tracks along the major groove of DNA (AT)_n_ (see **Fig 4**). Among eight key residues frequently in HB association with the Crick strand of DNA, the very front residue Arg135 (NH1/NH2) that initially bonds with A18 backbone (O2P; < 2 μs for configuration or config I), approaches forward first and forms HB with A19 (configure II at ∼2 μs), while other contacts almost remain intact (< 5 μs). At ∼5 μs, as the protein moves forward (state 3 in **Fig 3**), almost all contacts suddenly break within ∼ 20 ns (see **SI Fig S7A)**, except for Lys122-A16 and Lys144-A18 (config III); four of the front contacts (but not the one by Arg135) reform quickly (config IV, ∼ 30 ns), then the middle one (Arg131-A17, config V, ∼40 ns), the rear one (Lys125-A16, config VI, ∼60 ns), and finally, Arg135 forms contact with A19, which concludes the stepping cycle (config VII, ∼ 5.2 μs). During this stepping cycle for 1-bp, the protein oscillates back and forth and then moves forward (see **Fig 3**), tracking along the major groove. Further movements revealed in the simulation (5.2-10 μs) account for some sliding (∼2 bp) but not fully completed yet. It should be noted that during the sliding, individual contacts break in a way slightly differently (e.g. Tyr119 breaks contact first and then Arg135), while the middle and rear contacts break but have not yet reformed even though the COM of protein appears to move ∼ 2 bp (see **SI Fig S7B** for the detailed schematics of the contacts during sliding). The schematics of protein-DNA contacts on the Watson strand can be also found in **SI Fig 7C**): Though there are only 2-3 HBs formed occasionally, one can see that Arg118 breaks and reforms contact with the DNA backbone phosphates of T21’, T22’, T23’, and even T24’ throughout the 10-μs simulation (∼ 3 bp).

**Fig 4.**
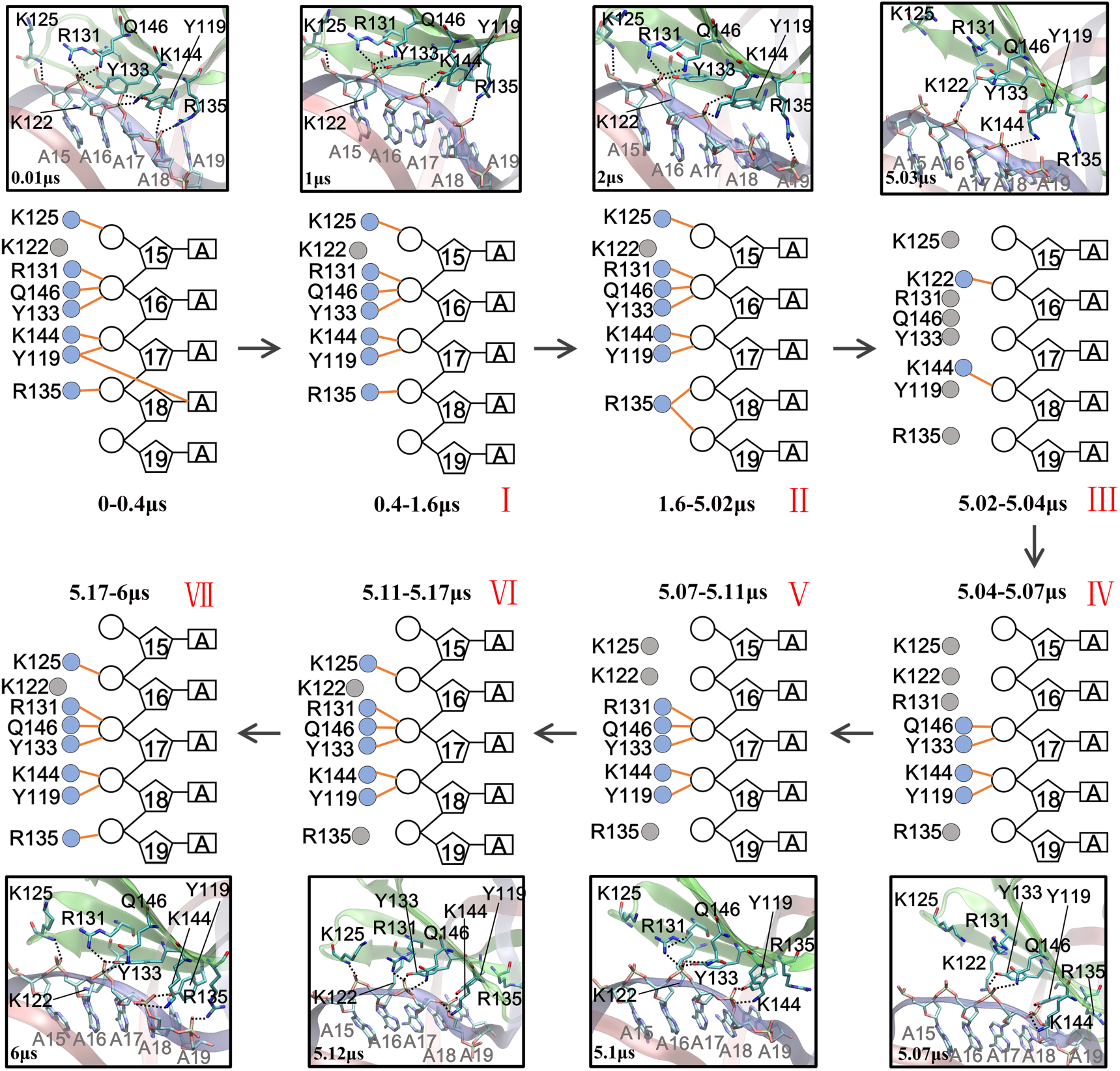
The stepping schematics and views of WRKY tracking forward along DNA during diffusion from the all-atom equilibrium MD. Since WRKY associates closely with the Crick strand of DNA, we show schematics of eight key residues (filled circle) from WRKY that make HB contacts with the Crick strand (open circle, pentagon and rectangle for the phosphate, sugar and base of a nucleotide, respectively). The HBs in the schematics are depicted in orange lines. The corresponding molecular views at the protein-DNA interface are illustrated (the Crick and Watson strand in blue and pink, respectively; WRKY in green).

Besides, we also calculated the protein interaction energetics with respective strands of DNA during the forward movements of WRKY on the DNA (see **SI Fig 7D**). During the 1-bp stepping stage (< 5 μs), the interaction of WRKY with the Crick strand (ele −137±32 and vdW −23±8 kcal/mol) does not distinguish that much from the Watson strand as in the previous specific or non-specific DNA binding case, though it remains much stronger than that with the Watson strand (ele: −52±26 and vdW −14±7 kcal/mol). The interaction between WRKY and the Watson strand indeed strengthens more than that in the specific binding case. Besides, the fluctuations of the WRKY interactions with DNA arise significantly for both DNA strands. The corresponding t-value and Pearson correlation coefficient in the electrostatic association become 101 and 0.07, with the t-value significantly reduced from that in the specific or non-specific binding case (see **SI Table S1**). Noticeably, during the sliding stage (> 5 μs), the interactions of WRKY with the Crick DNA strand (ele −110±35 and vdW −19±9 kcal/mol) and with the Watson strand (ele −71±32 and vdW −20±8 kcal/mol) become even less distinguishable and more correlated: The t-value and Pearson correlation coefficient in the electrostatic association change to 41 and −0.43, suggesting an increased similarity between the two-strand association energetics and an enhanced correlation between them, during the diffusion movements of protein and from stepping to sliding.

### Stepping backward and DNA strand crossing of WRKY to switch from major to minor groove DNA binding during diffusion

In the backward movement of WRKY along DNA, the protein also tracks along the major groove initially (< 6 μs; see **Fig 5**). It starts with Arg131 to the rear region squeezing on the peripheral Lys125 (from config I to II during 3-5 μs) to break both the back contact Lys125-A15 and a couple of front ones (config III, for ∼ 70 ns). After the middle contact region re-adjusts for a while (∼ 0.6 μs from IV to V), the initial set of contacts almost reform, except for the one from Lys125 to A14. However, WRKY seems to reduce its association then with the DNA (see **SI Fig S8A**), and crosses the Crick strand to move backward to the minor groove: One can find five residues (Arg131, Tyr133, Lys142, Lys144, and Gln146) res-establish their contacts with the Watson strand after crossing the Crick strand and slide for ∼ 2 bp backward along the DNA, with Lys142 and Lys144 associating with both strands upon the strand crossing (see detailed contact schematics and views for continue simulation between 6-10 μs in **SI Fig S8B**).

**Fig 5.**
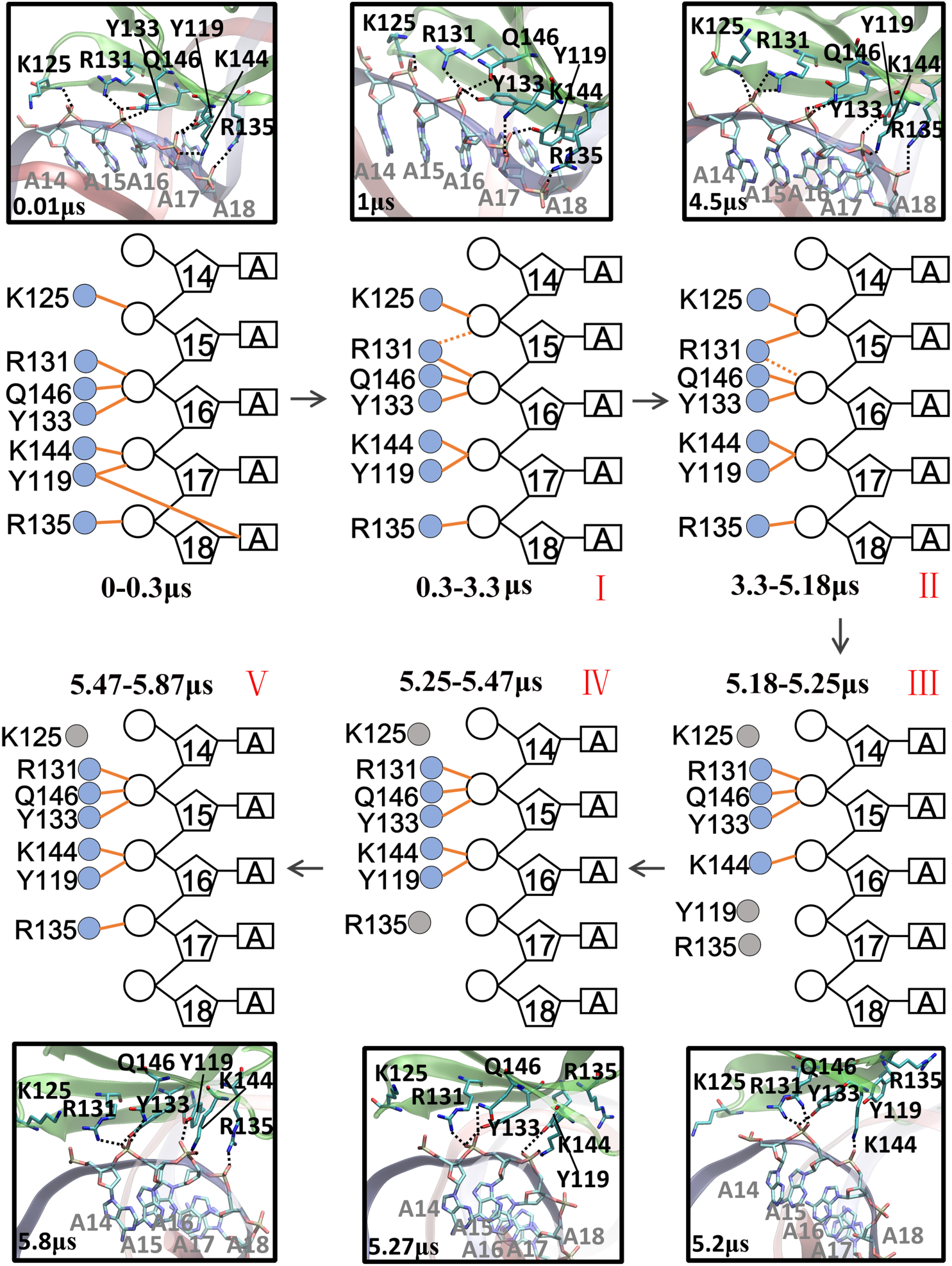
The stepping schematics and views of WRKY tracking backward along DNA during diffusion from the all-atom equilibrium MD simulation. We show schematics of seven key residues (filled circle) from WRKY that make HB contacts with the DNA Crick strand (open circle, pentagon and rectangle for the phosphate, sugar and base of a nucleotide, respectively). The HBs are also depicted in orange lines. The corresponding molecular views at the protein-DNA interface are also illustrated (the Crick and Watson strand in blue and pink, respectively; WRKY in green).

Similarly, we calculated for the backward movements of WRKY the protein-DNA interaction energetics, as for both the Crick and Watson strand (see **SI Fig 8C**). As in the forward direction during the 1-bp stepping (< 5 μs), the interaction of WRKY with the Crick strand (ele −144±28 and vdW −27±9 kcal/mol) does not appear quite different, on average, from the specific or non-specific binding case; the Crick strand association remains stronger than that with the Watson strand (ele: −56±29 and vdW −17±8 kcal/mol), until around the time the strand crossing happens (>5 μs). In particular, upon the strand crossing and further sliding, the WRKY interaction energetics with the two strands experience very large fluctuations (with the Crick strand ele −130±40 and vdW −20±9 kcal/mol; with the Watson strand ele −96±60 and vdW −20±14 kcal/mol) and become much less distinguishable than in all other cases (see **SI Table S1**). The corresponding t-values and Pearson correlation coefficients in the electrostatic association are 108 and 0.18 in the stepping stage (<5 μs), and become 24 and −0.04 in the strand-crossing and sliding stage (>5 μs). These two measures appear similar in the stepping stage for both forward and backward diffusive trajectories, while in the strand crossing case, the t value drops to a particularly low value as the two strand become almost “indistinguishable”, while the correlation remains quite low, in highly contrast with the forward sliding case, in which correlation is quite high (see **SI Table S1**).

## Discussion

In this work, based upon an obtained structures of DNA binding complexes of a plant TF, the WRKY protein domain, we demonstrated microseconds atomistic simulations of the essential diffusional motions of the TF along DNA with unprecedented structural dynamics details. In particular, the structural models of the WRKY-DNA complexes for both specific and non-specific DNA binding were constructed, together with a mutant protein K122A in complex with the originally specific DNA. The binding affinities of respective protein domain structures on the DNA were determined via ITC techniques. Our simulations then show that in all these protein-DNA binding complexes, one strand of DNA, the Crick strand, is bound preferentially. An onset of protein diffusion reveals for the WRKY domain protein COM in the non-specific DNA binding case (or for the mutant K122A), in which WRKY associates with the other strand, the Watson strand, via a different binding interface from that in the specific DNA binding case. Nevertheless, one could not yet detect movements of individual residues at the protein-DNA interface for the non-specific DNA binding complex (or for the mutant system).

With current high-performance computing technologies, one still could not sample the essential protein diffusion atomically for such a small TF attaching to an arbitrary non-specific DNA. Nevertheless, with the homogenous DNA (AT)_n_ adopted in our simulations, we were able to identify a complete 1-bp cycle of a full set of protein residues stepping on the DNA, followed by more or less stochastic motions. Indeed, by introducing the homogeneous DNA sequence with an exact 1-bp periodicity, we could sample comparatively high percentiles of 1-bp stepping motions (e.g. >40%, see **SI Fig S5**) for processive diffusion of WRKY along DNA in the coarse-grained or CG model of the protein-DNA complex, while the chance of the 1-bp stepping of WRKY lowers significantly for the random DNA sequence with a similar simulation set-up. In the CG representation, there is lack of protein side chains so that the HB interactions are excluded. Nevertheless, it appears that the DNA sequence periodicity (with the base identities partially preserved in the CG) impacts substantially on how the protein moves along DNA. The detailed features of the protein stepping and sliding along DNA during diffusion predicted in current work await experimental validations at base-bp resolution.

In the current atomistic simulations, both forward and backward movements of WRKY domain protein along DNA have been captured, as expected for the diffusive motions. Along the forward direction, we observe an elementary 1-bp stepping of WRKY, as individual protein-DNA contacts or HBs form on the DNA strand, break, and reform, following closely the major groove of DNA. Stochasticity is clearly noticeable right after the regular 1-bp stepping, as WRKY slides or steps in a size ∼ 2 bp, i.e, the individual contacts break and intend to reform after the 2-bp distance. Along the other or backward direction, WRKY steps similarly for 1-bp but backward, while the stochasticity shows more prominently when the protein crosses the Crick strand and moves onto the minor groove side of DNA. The types of diffusive motions of protein along DNA have been visualized in previous CG simulations lack of protein residue or side chain details (39, 40). Here we reveal the all-atom stepping or sliding motions of the TF protein, zoomed in with displacements of a full set of protein residues more or less synchronized at the DNA binding interface. The observations also make it easier to understand stepping behaviors of other nucleic acids walkers, or molecular walkers following a quasi-periodic track in general. For example, motor proteins such as DNA packaging motors or helicases have been detected with variable stepping sizes, e.g. 2-4 bps or larger, from single molecule measurements (41-45). Note that for those motor proteins, the stepping or sliding motions can be similarly and as fast as the TF proteins, but chemical catalysis or mechano-chemical coupling that supports directional movements of the motor proteins can be quite slow (e.g. milliseconds), so that they allow for sufficient experimental characterizations. Although various models could be proposed to explain those diverse stepping behaviors, from current observations from the WRKY domain protein simulation, one would infer that the multiple stepping sizes simply arise because of non-synchronized motions of individual protein residues with respect to the DNA binding interface, owing to variable DNA sequences and heterogeneous HB strengths. Besides, stochasticity plays a significant role in such stepping or sliding behaviors as the nanometer sized walkers relies on thermal fluctuations for coordinated motions.

Interestingly, we notice that even though WRKY distinguishes the two strands of DNA by preferentially binding to the Crick strand, particularly via electrostatic associations during the specific DNA binding, the biased association loosens somehow upon the non-specific DNA binding. The biased DNA strand association weakens highly significantly during diffusive movements of the protein, as the protein steps, slides, crosses the strand or switches the direction. That says, as the TF protein diffuses or searches along DNA, with the movements and stochasticity going on, the protein-DNA interactions become highly volatile, such that the preferential binding on the DNA strand cannot be stably maintained. However, when the TF reaches to regions with specific DNA sequences, the preferential electrostatic binding to particular DNA strand may possibly re-establish, so that further base scrutinization or recognition can be closely conducted, which further stabilizes the biased DNA strand association. We expect that with an integration of high-precision experimental detection with our developing simulation technologies, the energetic landscape and dynamics strategies of the TF protein searching on the genome would real more thoroughly.

## Materials and Methods

Detailed descriptions about obtaining the crystal structure, the setup of atomic and coarse-grained simulations, the Isothermal Titration Calorimetry (ITC) experiments, and the single molecule florescence experiments are provided in ***SI Appendix***.

## Acknowledgements

This work has been supported by NSFC Grant #11775016 and #11635002. JY has been supported by the CMCF of UCI via NSF DMS 1763272 and the Simons Foundation grant #594598 and start-up fund from UCI. We acknowledge the computational support from the Special Program for Applied Research on Super Computation of the NSFC Guangdong Joint Fund (the second phase) under Grant No. U1501501 and from the Beijing Computational Science Research Center (CSRC).

